# Design Guidelines for Dry Electrode Tip Geometry in Electroencephalography Measurements: A Proposal Based on the Scalp’s Mechanical Response

**DOI:** 10.1101/2025.07.30.667786

**Authors:** Shunya Araki, Shintaro Nakatani, Nozomu Araki

## Abstract

In electroencephalography (EEG) measurements using dry electrodes, a trade-off between signal stability and user comfort is a critical barrier to long-term, wearable applications. While various approaches have been proposed to address this issue, the mechanical impact of electrode tip geometry has not been adequately quantified, and most existing evaluations predominantly rely on subjective assessments. To address this gap with a quantitative, mechanics-based framework, the current study aimed to identify an optimized electrode tip geometry that minimizes mechanical stress on the scalp even under tilted contact conditions. Finite element analysis was conducted using strain energy density (SED)— a mechanical index known to correlate with neural impulse activity—as a quantitative indicator of the mechanical influence of tip geometry on the skin. Six types of electrode tip geometries, ranging from flat to hemispherical, were defined based on the ratio of fillet radius to prong radius. These geometries were analyzed under inclination angles from 0° to 5°, and their peak SED values were compared. Additionally, a geometry optimization using an iterative search algorithm was performed to minimize peak SED under the 5° tillt. The findings revealed that intermediate fillet geometries with gently rounded edges more effectively reduce peak SED under inclined conditions. Optimization further identified a geometry ratio of 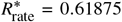 as the most effective tip geometry for minimizing mechanical loading under the specified conditions. These results offer a potential geometric design guideline for dry EEG electrodes that can help maintain user comfort across varying inclination angles.

## 1 Introduction

Electroencephalography (EEG) is a non-invasive and safe technique for recording brain activity. In recent years, EEG systems employing dry electrodes—which can considerably reduce application time compared to conventional electrodes—have garnered increasing attention [1]. Beyond neurological applications, the use of dry electrodes is now being explored in diverse fields such as longterm monitoring [2,3], communication [4], rehabilitation [5,6], and sports science [7,8]. However, current dry electrode EEG systems continue to face challenges related to user comfort [9]. In particular, the quality of contact between the electrode and the skin plays a critical role in balancing signal quality with wearer comfort.

Many dry electrodes used for EEG measurement employ a rigid, comb-like structure with multiple protrusions (prongs) designed to penetrate hair and establish stable contact with the scalp. These prongs exert pressure on the scalp to reduce skin–electrode impedance; however, insufficient pressure can raise impedance, thereby lowering the signal-to-noise ratio (SNR) of the EEG signal [10,11]. Conversely, excessive pressure compromises user comfort, making long-term EEG measurements challenging. Consequently, a trade-off arises between signal stability and wearer comfort [12]. To address this issue, previous studies have investigated the use of flexible electrode materials. Specifically, conductive silicone, polydimethylsiloxane, and polyurethane have been utilized to reduce mechanical stress at the electrode-skin interface [13–19]. These soft electrodes can be secured for prolonged use with head caps or medical tape [20]. Moreover, strategies have been proposed to incorporate elasticity into the electrode structure to reduce localized pressure and enhance wearer comfort [21– 24]. In parallel, previous studies have sought to enhance scalp contact by optimizing the geometry of prong tips. Accordingly, various designs have been proposed, including arch-shaped prongs that conform to the head’s curvature [25,26], tips with rounded or filleted features rather than cylindrical ones [21,27–29], and prongs with microstructured surfaces [30–32]. These designs are intended to resolve the trade-off between signal quality and user comfort. However, so far, their effectiveness has been evaluated primarily through subjective and sensory feedback assessments, andwhile quantitative studies examining the mechanical effects of prong tip geometry on the scalp remain scarce. Consequently, geometry optimization based on objective physical metrics has yet to be meaningfully advanced.

Given this background, the current study focuses on optimizing the tip geometry of rigid, multi-prong electrodes. Although previous studies have proposed a variety of geometries, quantitative evaluations of their mechanical effects on the scalp remain limited. To address this gap, the present study examines the impact of different tip geometries using a defined physical metric, excluding the consideration of flexible materials. Specifically, we examine the strain energy density (SED) generated within the skin owing to variations in electrode’s inclination angle, as illustrated in Fig. 1. Notably, SED describes the spatial distribution of energy stored in tissue and is known to correlate strongly with the intensity of mechanical stimuli perceived by mechanoreceptors and nociceptors [33,34]. Notably, pain models based on SED have been shown to replicate sensations assessed by the visual analog scale [35]. Accordingly, we use SED as an index to quantify the relative mechanical impact of prong tip geometry on the scalp. We hypothesize that the optimal geometry depends on the posture angle. To test this, we conduct finite element analysis on a single-prong model, treating the scalp as an isotropic linear elastic material while varying the posture angle, ψ. To ensure EEG signal stability, this analysis assumes a pressing force that exceeds a predefined threshold. By examining the resulting SED distributions —particularly their peak values— we assess the mechanical effects of each geometry under tilted contact conditions and to propose design guidelines for a geometry that minimizes mechanical scalp load.

**Fig. 1.**
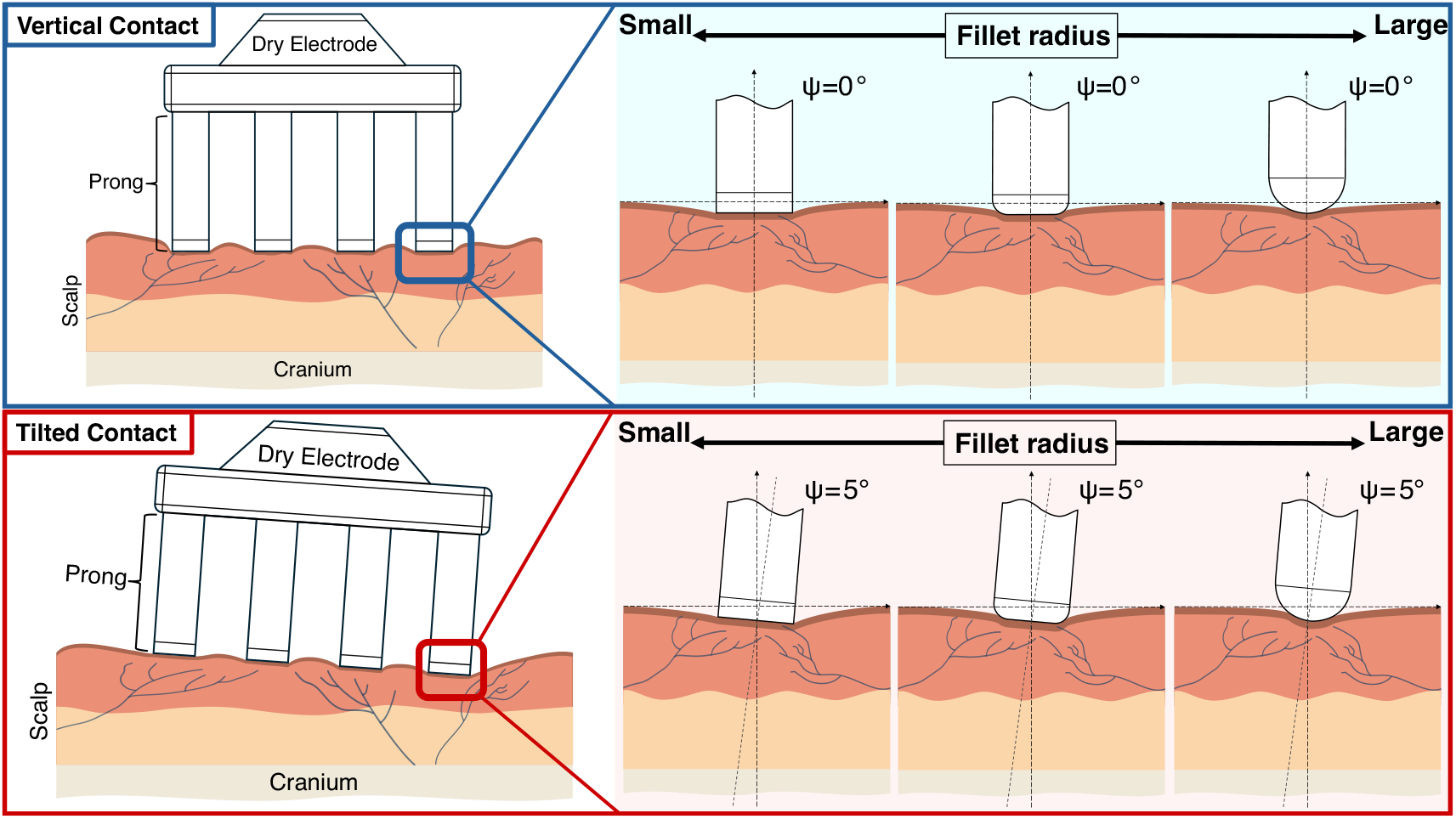
Conceptual diagram of the mechanical interaction between prong tip geometry and the scalp under vertical and tilted contact conditions.The figure illustrates how the electrode’s inclination angle (*ψ*) and prong tip geometry influence scalp deformation. The top row (blue frames) depicts axisymmetric contact under vertical pressing (*ψ* = 0°), while the bottom row (red frames) depicts non-axisymmetric contact under tilted pressing (*ψ* = 5°). As the inclination angle increases, scalp deformation becomes more pronounced. The nature of the contact interaction also varies with prong tip geometry, with each geometry expected to induce characteristic deformation patterns and stress concentrations, especially under tilted conditions.

## 2 Electrode Prong–Scalp Contact Model

### 2.1 Scalp Modeling

This section describes the scalp model used in this study by outlining the relationship between mechanoreceptors, the anatomical and mechanical structure of the scalp, and the mechanical properties of the scalp.

#### 2.1.1 Mechanoreceptors and Mechanical Stimuli

Hairy skin features low-threshold mechanoreceptors (LTMRs), which detect innocuous mechanical stimuli near hair follicles, and high-threshold mechanoreceptors (HTMRs), which respond to noxious stimuli [39]. Among these, HTMRs are further classified into AHTMRs, which respond to rapid mechanical input, and C-HTMRs, which react to slow, sustained stimuli [40]. These receptors convert mechanical stimuli into neural impulses that are transmitted to the brain and interpreted as pain. Previous studies have reported a strong correlation between the firing frequency of LTMRs and the SED within the skin [33,41]. Furthermore, in a model developed to represent the pain response mediated by HTMRs, neural activation occurs when SED exceeds a specific threshold, indicating that SED may serve as a valid indicator of neural activity [35].

Based on these findings, SED is regarded as a reliable physical metric for representing the intensity of neural activation. The activity of both LTMRs and HTMRs is thought to correlate with SED through a shared mechanical-to-electrical energy conversion mechanism [34]. Moreover, the validity of SED as an evaluation metric for HTMR activity is strongly supported by proposed pain models that emphasize biomechanical energy within the skin. However, the present analysis does not verify the actual activation of HTMRs. Accordingly, SED is not used as a direct indicator of nociceptor activity. Instead, it serves solely as a physical index for quantitatively comparing the relative mechanical impact exerted by different prong tip geometries on the scalp.

Given that dry electrodes predominantly apply static pressure to the scalp during EEG measurements, using such an index to compare contact geometries provides a meaningful basis for identifying structural factors that may underlie differences in pain perception.

#### 2.1.2 Anatomical Structure of the Scal

As depicted in Fig. 2(a), the human scalp comprises five distinct fibrous tissue layers [36–38]. The outermost layer is the *skin*, which includes the epidermis and dermis; this layer is the thickest and contains numerous hair follicles. The skin’s mechanical behavior is primarily governed by the dermis, and mechanoreceptive nerve endings are distributed around the hair follicles [42]. Beneath the skin is the *connective tissue* layer, which is rich in adipose tissue and densely, supplied with blood vessels. This is followed by the *aponeurosis* (also known as the galea), a thin, dense, and resilient fibrous layer. The fourth layer is the *loose areolar tissue*, characterized by a relatively sparse vascular network. The deepest layer is the *pericranium*, a membranous tissue that tightly adheres to the surface of the skull. Notably, scalp thickness varies considerably depending on factors such as age, sex, body mass index, and anatomical location. Reported values range from approximately 2.0–4.0 mm in the frontal region [43,44], 2.2–5.0 mm in the parietal region [45], 6.2–11 mm in the temporal region [46], and 5.0–11 mm in the occipital region [47].

**Fig. 2.**
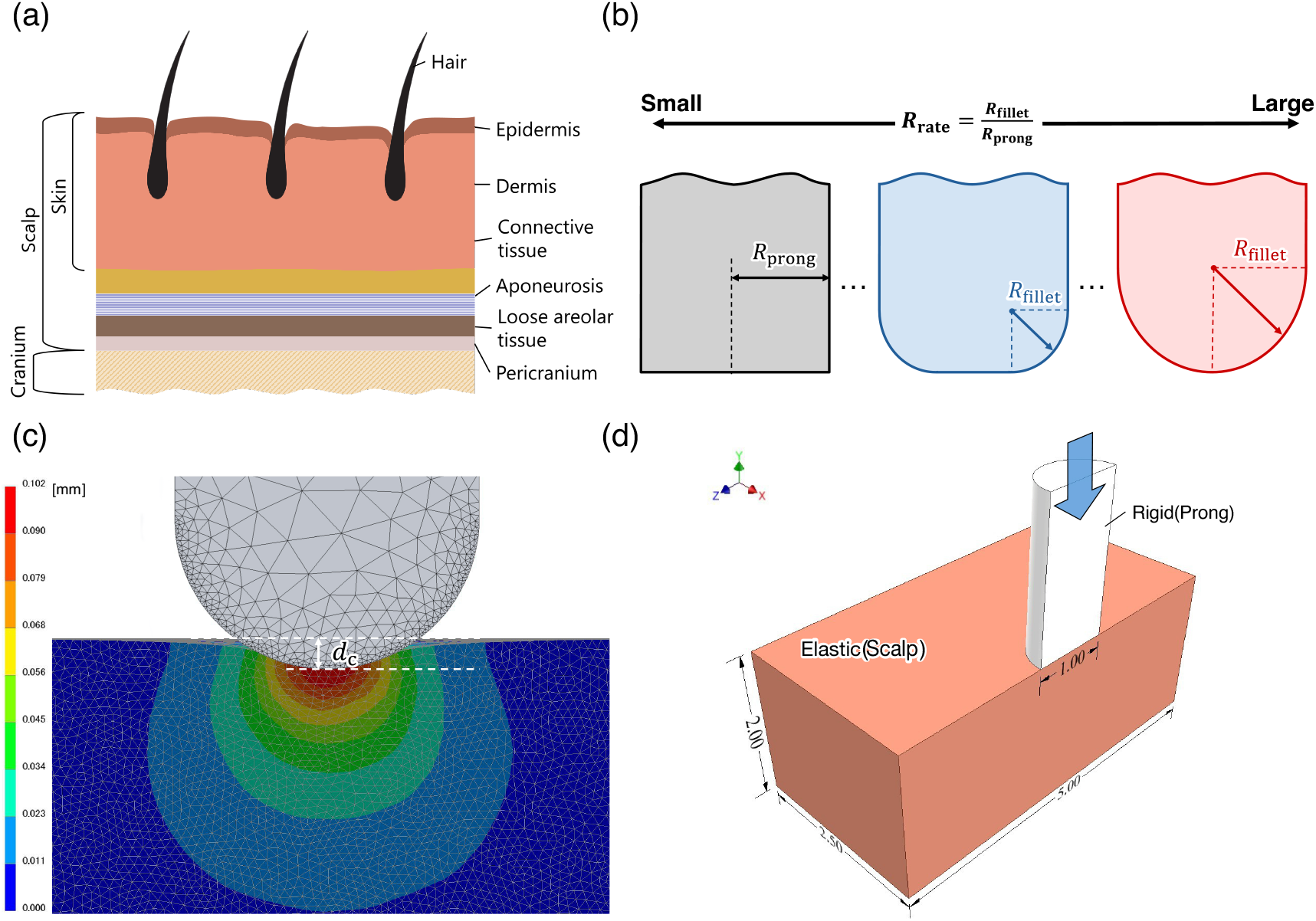
Overview of the anatomical structure and FEM modeling framework considered in this study: (a) schematic representation of the scalp’s layered structure, from the skin (epidermis and dermis) to the cranium [36–38]; (b) definition of prong tip geometry, quantified by the shape ratio *R*_rate_ = *R*_fillet_ *R*_prong_; (c) results of a preliminary analysis to determine the common indentation depth (*d*_*c*_ = 0.1 mm) used for all geometries, based on the minimum pressing force required for EEG signal stability [18]; (d) three-dimensional finite element model of prong–scalp contact, where the prong (white) is modeled as a rigid body and the scalp (orange) as an elastic body.

The objective of this study is not to replicate the detailed layered structure or physiological responses of the skin with high fidelity but to assess the mechanical influence of different prong tip geometries. Accordingly, the scalp was modeled as a homogeneous elastic body with a uniform thickness of 2 mm, based on two primary considerations. First, this approach ensures consistency with a previous study [29] wherein prong tip design was optimized using FE analysis based on von Mises stress, with subjective comfort validated experimentally. Notably, von Mises stress and SED are both indicators of stress concentration derived from principal stresses and have been demonstrated to correlate strongly when used to compare localized mechanical stimulus distributions. Second, while multilayer models capable of accurately reproducing the skin’s layered architecture are appropriate and essential for simulations of phenomena such as wrinkle formation or wound healing —where interlayer differences in stress and deformation are critical— they are not necessary for evaluating mechanical responses associated with geometry and contact conditions. Studies on vacuum-assisted delivery and dry electrode design have indicated that a homogeneous skin model is sufficiently accurate for such applications [48,49]. In line with these findings, the present study adopts a homogeneous scalp model, which offers a reasonable and reproducible basis for comparing stress distributions across different prong tip geometries.

#### 2.1.3 Mechanical Properties

The skin exhibits various mechanical behaviors, including anisotropy, nonlinearity, and viscoelasticity [50]. Although several constitutive models have been proposed to capture these characteristics, computational approaches specifically designed to simulate the mechanical response of the scalp remain limited. Existing frameworks include a hyperelastic membrane model based on the study of Tong and Fung [51,52], a two-layer model incorporating the first-order Ogden formulation [53], and a simplified single-layer linear elastic model [54]. In parallel, studies examining the relationship between neural responses and mechanical indices have reported a strong correlation between Merkel cell firing rates and SED [33]. Although these studies focus primarily on the fingertip, the structural and sensory receptor similarities between fingertip and scalp tissue suggest that SED likewise serves as a valid indicator of mechanical effects for the scalp. Building on these insights, the current study assesses the mechanical response of the scalp using SED as a mechanical indicator of neural activation. Within this framework, the skin is modeled as a linear elastic body. he objective is not to replicate large-scale surface deformations or obtain absolute response values with high precision but to capture relative differences in mechanical behavior across prong tip geometries and variations in contact conditions. This modeling approach is supported by previous research demonstrating that linear elastic models can yield meaningful insights for such comparative analysis [33,41].

To represent the skin as a linear elastic material, two parameters must be defined: Young’s modulus and Poisson’s ratio. Among these, Young’s modulus, which characterizes the skin’s mechanical behavior, has been measured using a variety of experimental techniques, including indentation, tensile, torsion, and suction tests [55]. These tests have yielded the following range of values: 19.1–22.31 MPa (tensile tests [56]), 1.09–651 kPa(indentation tests [57]), 0.02–0.1 MPa(torsion tests [58]), and 130–260 kPa(suction tests [59]). Given that the dermis is the primary contributor to the skin’s mechanical response, a Young’s modulus of 0.066 MPawas adopted in this study, based on the inverse FE analysis results reported by Hara et al [60]. This value falls within the range obtained from indentation tests and is considered appropriate for the purposes of this study. Additionally, following the modeling assumptions of Srinivasan et al. [33] and Sato et al. [41], the skin is assumed to be isotropic and nearly incompressible. Although a perfectly incompressible material has a theoretical Poisson’s ratio of 0.5, most finite element software, including Autodesk Inventor Nastran, cannot accommodate this value. Hence, a Poisson’s ratio of 0.48 was adopted in this study. This value closely approximates the theoretical upper limit and has been demonstrated to effectively reproduce incompressible behavior [61–63].

### 2.2 Prong Structure

Rigid dry electrodes, which are the focus of this study, typically feature a comb-like, multi-prong structure with several protrusions extending from the base. The choice of this configuration is primarily determined by the electrical requirements of dry electrodes. To ensure a stable electrical connection, dry electrodes must pass through the hair layer and allow the prong tips to establish direct contact with the scalp.

However, human scalp hair is typically dense. A study on Arab males aged 18–30 reported hair densities of 147.9 ± 3.7 hairs / cm^2^ in the frontal region, 152.3 ± 6.3 hairs / cm^2^ in the temporal region, and 158. ± 1 6.7 hairs / cm^2^ in the occipital region [64]. To navigate such densely distributed hair and reach the scalp, individual prongs must have a diameter small enough to pass between hair shafts. For instance, based on the occipital region’s hair density, approximately 1.5 hair strands would be present within a 1 mm × 1 mm area where a 1-mm-diameter prong is applied. This suggests that a prong of this size can likely pass through hair gaps and reach the skin surface. Accordingly, this study adopts a prong radius of 0.5 mm, which is considered appropriate for insertion through hair.

Previous research has revealed that prong diameter, prong count, and tip geometry must be comprehensively designed to improve the SNR of an EEG signal. For example, one study evaluated the optimal number of prongs required to balance EEG signal stability with user comfort, assuming a prong radius of 0.5 mm and a hemispherical tip [18]. Notably, among the abovementioned factors, tip geometry plays a particularly important role in optimizing electrode–scalp contact. Reported designs for tip geometry include a dry electrode featuring five semicircular arches arranged in parallel at 1 mm intervals and a reverse-curve, arch-shaped structure that conforms to the head’s curvature to enhance scalp contact [25,26].

Many current commercially available dry electrodes employ a multi-prong structure, often featuring filleted tip designs [65,66]. Compared to flat-tipped designs, such filleted prongs are expected to alleviate stress concentration at the edges, thereby reducing the mechanical load on the scalp. Building on this background, the present study focuses on prong tip geometry. As illustrated in Fig. 2(b), we define six tip geometries that represent a continuous transition from a flat to a hemispherical shape. These geometries are defined by setting the ratio *R*_rate_ to 0.0, 0.2, 0.4, 0.6, 0.8, and 1.0, based on Eq. 1. Subsequently, the SED is evaluated for each of these ratios.

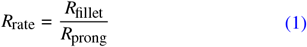

where *R*_fillet_ denotes the fillet radius at the prong tip, and *R*_prong_ represents the prong radius.

For the contact model considered in this study, we evaluate a single prong located at the periphery of a multi-prong electrode structure. This approach is based on the assumption that a peripheral prong serves as the most conservative evaluation case, as localized stress concentrations and poor contact are most likely to occur in this region. Moreover, to ensure structural and numerical simplicity, a single-prong model offers a valid method for evaluating tip geometry during the initial design stage.

### 2.3 Modeling of Contact Conditions

#### 2.3.1 Indentation Depth

To enable a valid comparison of prong tip geometry effects under various contact conditions, the stability of EEG signal acquisition must be ensured. A key factor influencing EEG signal quality is the skin–electrode impedance (*Z*_se_). Given the trade-off between comfort and signal quality, an appropriate pressing force must be applied to maintain *Z*_se_ below a certain threshold (*Z*_se,max_). Previous studies have reported that EEG signal quality is maintained when *Z*_se_ < 350 kΩ [67].

Based on this value, we set the minimum pressing force to *F*_min_ = 0.2 N, following the approach of Fiedler et al. [18], who reported that a dry electrode with 30 hemispherical prongs achieved a *Z*_se_value of348 kΩ under this force. Accordingly, the pressing force per prong is *F*_min_ = 0.0066 N. Rather than using force as a common condition, we selected indentation depth *d*_c_—measured when a hemispherical prong is vertically pressed with this minimum force—as the unified condition across all geometries. To determine this value, we performed an analysis and observed that applying *F*_min_ = 0.0066 N yields an indentation depth of *d*_*c*_ = 0.1 mm. This approach enables a clearer comparison of the contact states produced by different tip geometries. Given that indentation depth directly reflects geometric deformation, it plays a critical role in evaluating local strain and SED distributions across the contact region. Moreover, it facilitates quantitative assessment of contact characteristics such as contact area and stress concentration.

In contrast, applying a fixed pressing force across all geometries would lead to inconsistent deformation levels. Consequently, variations in contact behavior or SED distribution would become difficult to attribute to either the force magnitude or the geometry itself. To resolve this ambiguity, we adopted indentation depth *d*_c_ —defined as the indentation depth of a hemispherical tip— as a standardized condition for isolating the effect of tip geometry on contact mechanics. The baseline indentation *d*_c_ is illustrated in Fig. 2 (c).

#### 2.3.2 Modeling of Tilted Contact Scenarios

Non-ideal contact states in EEG measurements using dry electrodes primarily stem from two factors. First, the human head is not a perfect sphere but rather an ellipsoid with inter-individual variability. According to the cephalic index—a commonly used head geometry metric calculated as the ratio of head width to length—head shapes are typically classified as brachycephalic (> 80%), mesocephalic (75–80%), or dolichocephalic (< 75%) [68]. Additionally, studies based on three-dimensional models reconstructed from computed tomography scans have reported substantial surface roughness [69]. Second, the fixation method strongly influences the contact state. While rigid headgear can help maintain proper electrode orientation to some extent, the flexible fabric caps widely favored for their comfort and adaptability may fail to ensure that the pressing force is applied perpendicularly to the scalp. Consequently, these factors can introduce a tilt in the electrode’s postural angle (ψ) —defined as the angle between the vertical direction and the electrode axis— leading to a non-ideal contact state.

The mechanical conditions that lead to this inclination angle (ψ) can be grouped into three categories. The first, initially-inclined pressing, arises when local scalp irregularities cause the electrode to tilt at the moment of placement. The second, rotation-induced contact, results from fixation methods —such as flexible caps— that fail to maintain a normal pressing force. This deviation alters the force vector and induces a rotational moment. The third, disturbance-induced tilt, stems from influences such as head movement during measurement.

Among these conditions, the present study focuses primarily on rotation-induced contact. Because previous studies have highlighted the critical importance of maintaining normal-direction contact pressure to ensure user comfort, rotation-induced contact, which inherently deviates from this ideal, becomes a central issue [18]. Moreover, this scenario frequently occurs in practical settings involving flexible caps and is well suited for systematic comparison, as it can be reliably reproduced in static FEM analysis. In addition to rotation-induced contact, initially inclined pressing represents another important case, as it stems from head geometry. An additional analysis was thus conducted for this scenario, and the corresponding results and discussion are provided in the Supplementary Material.

In this study, the inclination angle ψ was varied from 0° to 5°. A central objective was to systematically evaluate how the tip fillet affects the mechanical response during tilted contact, which requires consistent conditions for fair comparison. he flat geometry (*R*_rate_ = 0.0), which lacks a fillet, was designated as the baseline geometry for assessing geometric effects. The upper limit of 5° was selected because it represents the maximum inclination angle at which the baseline geometry maintains full contact with the scalp surface, without edge lift-off. Evaluating all geometries within this range allows a direct comparison of their effects on SED distribution under identical surface contact conditions.

## 3 Analysis

This chapter details the methodology of the FEM analysis used to quantitatively evaluate the mechanical influence of different electrode tip geometries on the scalp. It is organized into three parts. First, it describes the development of the three-dimensional model used as the basis for the analysis, along with the simulation setup. Second, it outlines the methods for acquiring and processing data from the simulation. Finally, it presents the evaluation metric used to assess the performance of the geometries based on the simulation results.

### 3.1 FEM Framework and Condition Setup

In this study, a three-dimensional FE model was developed and is illustrated in Fig. 2(d). A nonlinear static analysis was then conducted using Autodesk Inventor Nastran 2025 [70]. The model was meshed using tetrahedral elements, with a finer mesh applied near the prong tip and the contact region of the elastic body. The mesh size ranged from 0.01–0.2 mm, resulting in approximately 160,000-200,000 elements. The coordinate system was defined with its origin at the initial contact center between the prong and elastic body. The indentation direction was set as the Y-axis, while the horizontal direction parallel to the contact surface was defined as the Z-axis. All nodes on the skull-side surface of the elastic body and on the symmetry plane were fixed. The contact between the prong and the elastic body was defined as frictionless and no-slip. The prong tip was modeled as a rigid body, while the elastic body was represented as deformable. Both geometric and contact nonlinearities were considered in the analysis.

The inclination angle ψ (defined as the angle between the electrode axis and vertical direction) was varied from 0° to 5° in the analysis. For each condition, a displacement-controlled boundary condition was applied to maintain a constant indentation depth *d*_c_ at the center of the prong tip.

### 3.2 Data Acquisition and Processing

Our analysis focused on a vertical cross-section (Y–Z plane) through the center of the contact region in the elastic body, where mechanical responses are most pronounced. Within this plane, a 0.5 mm × 0.5 mm region was selected, containing approximately 2,200 tetrahedral elements. To ensure consistency across all simulation conditions, evaluation points were defined at identical spatial coordinates for all prong geometries and inclination angles. Notably, the obtained SED data were susceptible to local noise owing to mesh discretization, numerical integration errors, and minor variations in element size. These noise sources could obscure the assessment of geometry-dependent trends. To mitigate this, statistical processing was applied to exclude outliers prior to analysis. Specifically, a 95% confidence interval (CI) was used for the 2,200 SED values, with data below the 2.5 percentile and above the 97.5 percentile treated as outliers and excluded. This process removed extreme or erroneous values and improved the reliability of the SED data. The same statistical filtering was applied across all conditions to ensure a fair and consistent basis for comparison. After outlier removal, approximately 2,000 valid SED values remained per case. This number was considered sufficient for robust analysis.

### 3.3 Evaluation

This study uses SED as a physical index to quantitatively assess the relative mechanical impact of various electrode tip geometries on the scalp. Drawing on prior research suggesting a link between SED and the response of mechanoreceptors in the skin, this parameter is adopted here as an objective metric for comparing the intensity of mechanical stimuli potentially related to comfort.

SED (*U*_0_), which quantifies the energy stored per unit volume as a material deforms, is defined as in Eq. (2).

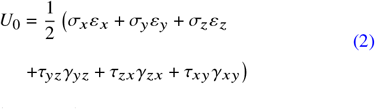

Here, σ_*i*_ and ε_*i*_(*i* = *x, y, z*)denote the normal stresses and strains, respectively, while τ_*j k*_ and γ_*j k*_ (*j k* = *yz, zx, xy*) represent the corresponding shear stresses and strains.

The primary evaluation metric in our analysis is the maximum SED (SED_max_), as it is considered most directly associated with the onset of discomfort or pain. To minimize or exclude the influence of computational noise, SED_max_ is defined as the highest value within the 95% CI of all evaluated elements. Focusing on this single metric enables a direct comparison of the most extreme mechanical stimulus generated by each geometry and its associated risk.

The following chapters present the analysis results for each geometry based on SED_max_, along with a quantitative comparison and discussion of their performance.

## 4 Results

### 4.1 SED_max_ Responses for Each Geometry Ratio

As illustrated in Fig. 3 (a), under vertical contact conditions, SED_max_ increases with the tip geometry ratio. A particularly sharp rise occurs when the ratio exceeds approximately *R*_rate_ = 0.6. In contrast, under tilted contact conditions, SED_max_ follows a concave-upward trend, reaching a minimum at 0.6. This is an important finding, suggesting that contact area distribution and deformation absorption may be optimized at a specific geometry ratio. Furthermore, the increase in SED from 0° to 5° is most pronounced at smaller geometry ratios, with the largest change observed at *R*_rate_ = 0.0. This indicates that for flat geometries, even slight tilting causes strong localized deformation, increasing SED_max_. Moreover, the 0° and 5° plots in Fig. 3 (a) do not intersect, indicating that SED_max_ consistently increases under inclined conditions across all geometry ratios.s

**Fig. 3.**
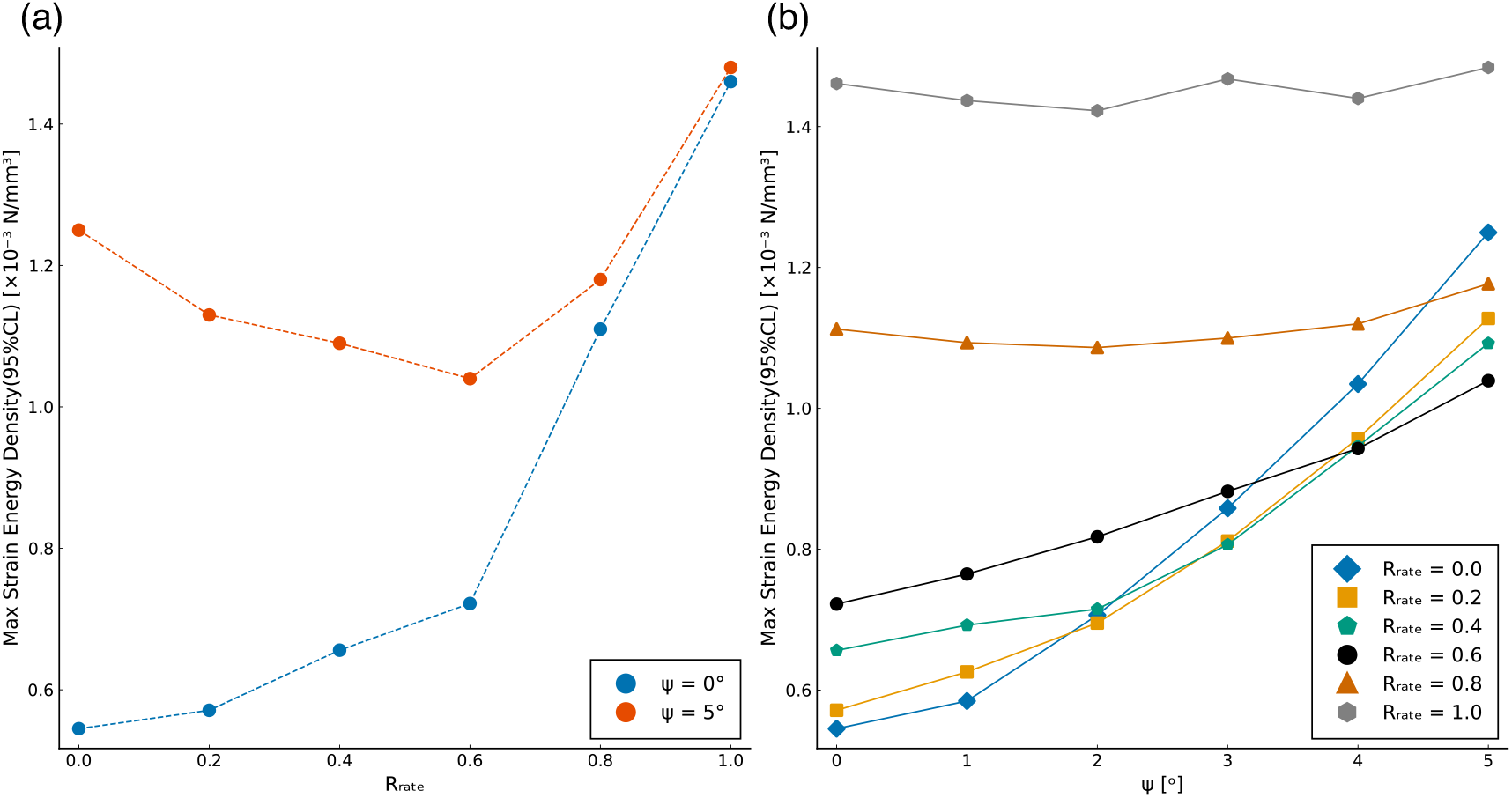
Relationship between prong tip geometry, inclination angle (*ψ*), and maximum SED (SED_max_): (a) SED_max_ as a function of the geometry ratio (*R*_rate_) under two conditions—vertical contact (*ψ* = 0°) and tilted contact (*ψ* = 5°). (b) SED_max_ as a function of the inclination angle (*ψ*) for six tip geometries.

### 4.2 Changes in SED_max_ with Varying Inclination Angles for Each Geometry

As depicted in Fig. 3 (b), the inclination angle (ψ = 0° – 5°) was incrementally varied across six tip geometries, and the resulting trends in SED_max_ were compared. The results revealed an increase in SED_max_ for all geometries under tilted conditions, although the rate of increase and initial values varied with geometry. In particular, geometries with*R*_rate_ values of 0.4 and 0.6 exhibited relatively small increases in SED_max_ as inclination increased, demonstrating more gradual responses than the other geometries. In contrast, the flat (*R*_rate_ = 0.0) and hemispherical (*R*_rate_ = 1.0) geometries exhibited distinct behaviors. Specifically, for (*R*_rate_ = 0.0, SED_max_ increased markedly with increasing inclination, especially beyond 3_°_. Meanwhile, for *R*_rate_ = 1.0, SED_max_ remained nearly constant, indicating that the tilt-induced variation in stress response was minimized.

Figure 4 illustrates SED distributions for three representative geometries (*R*_rate_ = 0.0, 0.6, and 1.0) at ψ = 0° and 5°, highlighting differences in spatial distribution. For the flat geometry (*R*_rate_ = 0.0), inclined contact caused the SED to become highly localized at the contact edge, resulting in an asymmetric distribution. In contrast, the intermediate geometry (*R*_rate_ = 0.6)exhibited no strong localization, and the SED distribution remained relatively smooth and widespread. For the hemispherical geometry (*R*_rate_ = 1.0), the SED was evenly distributed across the contact area, and the effects of inclination on the distribution pattern were minimal. However, across all cases, high SED consistently concentrated just below the center of the contact area (*y* ≈ − 0.1mm), indicating a structural tendency for deep stress concentration. These results suggest that while the hemispherical geometry offers the most stable response to inclination, it also tends to sustain a consistently high-stress state.

**Fig. 4.**
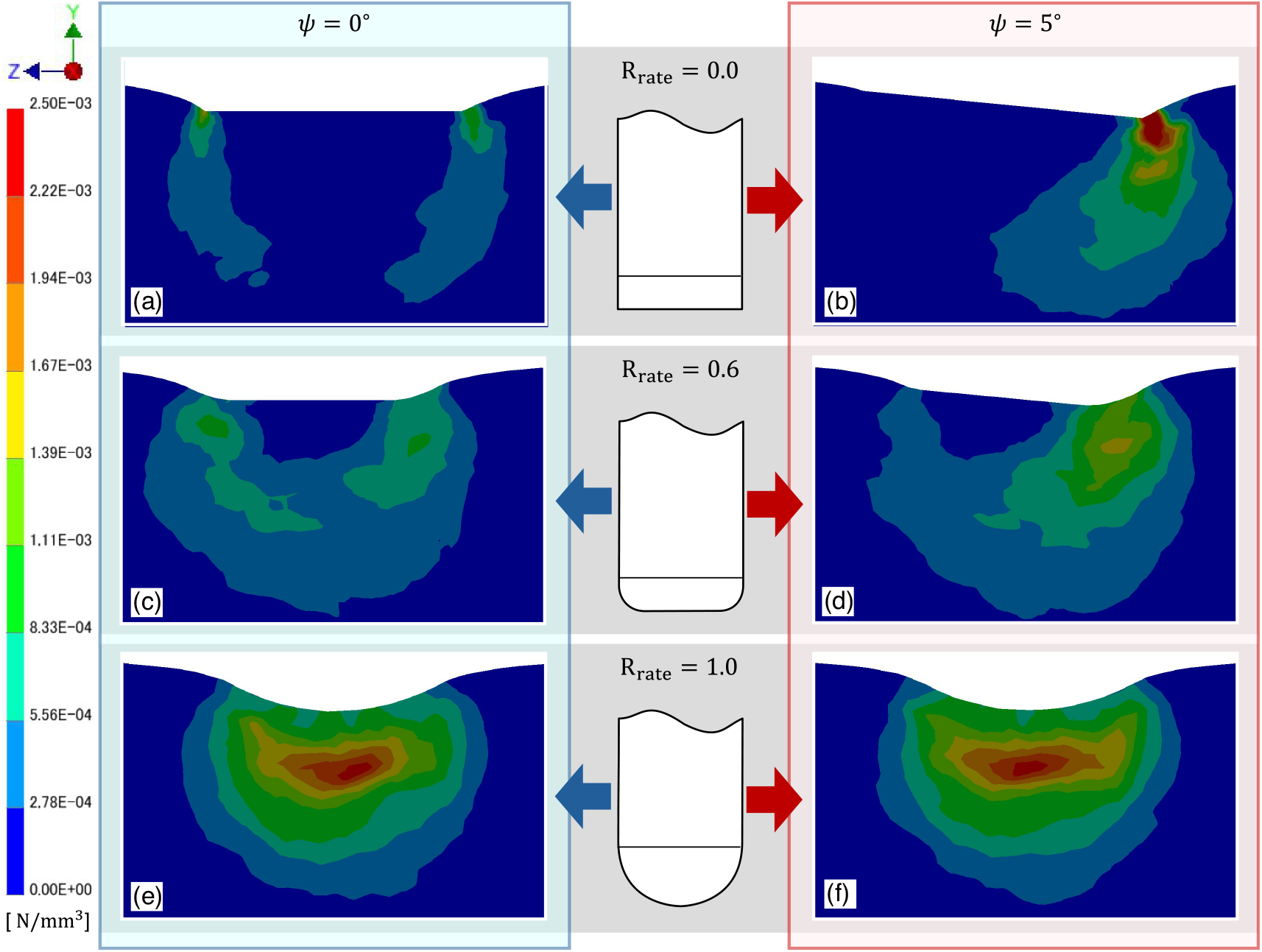
Comparison of SED distributions for representative tip geometries and inclination angles. Heatmaps display the SED distributions for three geometry ratios (*R*_rate_ = 0.0, 0.6, and 1.0). Two contact conditions—vertical (*ψ* = 0°) and tilted (*ψ* = 5°)— are presented side by side for comparison. Panels (a) and (b) correspond to *R*_rate_ = 0.0, (c) and (d) to *R*_rate_ = 0.6, and (e) and (f) to *R*_rate_ = 1.0.

## 5 Investigation of Optimal Geometry Based on SED_max_ Minimization

### 5.1 Optimization Problem Formulation

To maximize electrode comfort, we formulated a geometry optimization problem aimed at minimizing SED_max_ in the skin, using the tip geometry ratio (*R*_rate_) as the design variable. The objective function was defined as the worst-case mechanical stimulus within the evaluated range of inclination angles, 0°≤ ψ ≤5°. The optimization problem is thus expressed as in Eq. (3).

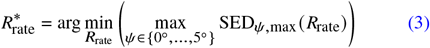

As depicted in Fig. 3 (b), the objective function value was consistently determined by the ψ = 5° condition. Accordingly, *f* (*R*_rate_)= SED_max_ (*R*_rate_, ψ = 5°) was adopted as the objective function for the subsequent search. Furthermore, as illustrated in Fig. 3 (a), the function displays a downwardly convex profile, enabling the use of an iterative search algorithm to efficiently identify its minimum.

### 5.2 Exploration Algorithm and Optimization Results

An iterative search algorithm was used to optimize the geometry, with the search direction determined based on comparisons between adjacent evaluated points. The search process was initialized at a geometry ratio of *R*_rate_ = 0.6, identified in preliminary evaluations as a favorable starting point. This value was treated as the current minimum, denoted by 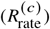, with an initial learning rate (step size) set to η = 0.1.

At each iteration, the next candidate value, 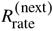, was computed by comparing the objective function at 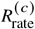 with that at a reference point, 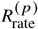—typically the value assessed in the previous step— according to Eq. 4.

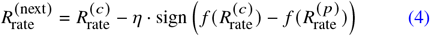

An additional analysis was then performed to evaluate the objective function at 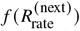. If the result improved upon the current minimum 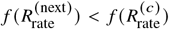, the current minimum was updated to 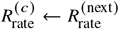, and the learning rate was maintained. If no improvement occurred and the convergence criterion was not met, the learning rate was reduced using a decay rate of *D*_rate_ = 0.5.

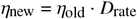

The search was terminated when improvements in the objective function became adequately small. Specifically, the search concluded when the gradient, approximated by the difference between the objective function values at the candidate point and the current minimum divided by the step size, met the following criterion.

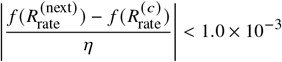

Here, *f* ^(*i*)^ denotes the objective function value at the current iteration, *f* ^(min)^ represents the minimum value observed so far, and η_old_ signifies the learning rate used in that step.

Following this procedure, the search converged after 13 additional evaluations, yielding the following optimal geometry ratio:

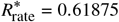

The corresponding objective function value at this point was 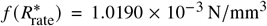. As illustrated in Fig. 5, the search process confirmed that a well-defined local minimum was successfully identified among the tested configurations.

**Fig. 5.**
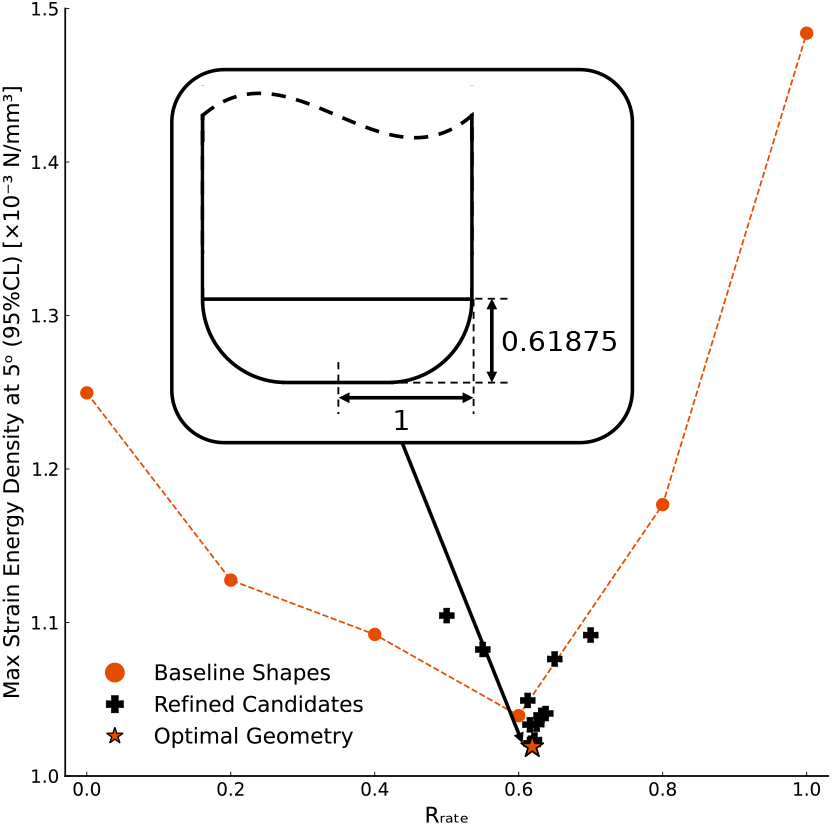
Optimization process via iterative search to minimize SED_max_ under tilted contact (*ψ* = 5°). The figure presents the objective function, SED_max_ at *ψ* = 5°, plotted with respect to the geometry ratio, *R*_rate_. The iterative search was initiated from baseline shapes (orange circles), evaluated additional candidates (black crosses), and converged to the optimal geometry (orange star) at 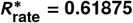.

### 5.3 Comparative Analysis of the Optimal Geometry

A detailed analysis was conducted to evaluate how the identified optimal geometry 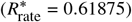 responds to changes in inclination angle compared to other representative geometries. As illustrated in Fig. 6, although this optimal geometry displays slightly higher SED_max_ than the flat geometry (*R*_rate_ = 0.0) under normal contact (ψ = 0°), the rate of increase in SED_max_ with inclination angle is more gradual. Moreover, across the full range of inclination angles, in contrast to the hemispherical geometry (*R*_rate_ = 1.0), which consistently exhibits high stress concentrations, the optimal geometry avoids sustained stress elevation and tends to maintain lower SED_max_ values.

**Fig. 6.**
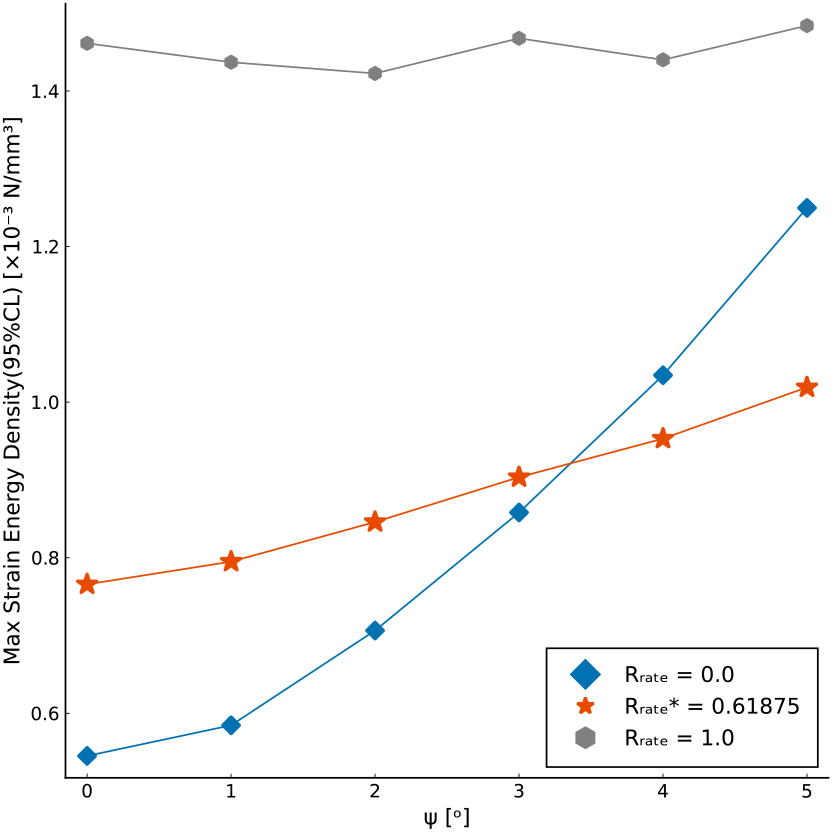
Comparative response of SED_max_ to inclination angle for the optimal and representative tip geometries. The figure illustrates the SED_max_ response to inclination angle, (*ψ*), for three characteristic tip geometries: flat (*R*_rate_ = 0.0), optimal 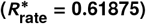, and hemispherical (*R*_rate_ = 1.0). While the optimal geometry exhibits slightly higher SED_max_ than the flat geometry under normal contact (*ψ* = 0°), it responds more gradually to increasing inclination angle and consistently maintains lower values than the hemispherical geometry.

## 6 Discussion

### 6.1 Validity of the Optimal Geometry and Comparison with Mainstream Designs

The goal of this study was to identify an electrode tip geometry capable of maintaining user comfort under tilted contact conditions, which are common in practical settings. Our analysis quantitatively demonstrated that both the mainstream hemispherical geometry (*R*_rate_ = 1.0) and flat geometry (*R*_rate_ = 0.0), representing the two extremes, offer distinct advantages and limitations.

The hemispherical geometry, widely used in commercial products, appears particularly effective in maintaining stable skin–electrode impedance, as its contact area remains relatively consistent with changes in inclination angle [15,16,22,23,27,29,65, 66]. This low sensitivity to inclination was also confirmed in our analysis, which revealed that the hemispherical geometry exhibited the smallest variation in SED_max_ with respect to tilt.

However, a key finding of this study is that the observed ow sensitivity to tilt occurs at a high absolute value, reflecting a previously under-addressed issue: the continuous application of a high mechanical load to the scalp. This persistent stress can be a direct source of discomfort and pain, particularly in situations where the pressing force is not well controlled, such as in daily wearable use. This suggests that comfort may be compromised in favor of impedance stability.

In contrast, the optimal geometry 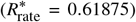 identified through our iterative search offers a clear solution to this issue. As illustrated in Fig. 6, unlike the hemispherical geometry, which consistently maintains a high SED_max_, the optimal geometry keeps the value low under normal contact and limits its increase as the inclination angle rises. This well-balanced behavior addresses the main limitation of the hemispherical geometry—its high absolute value—while retaining some of its low sensitivity to inclination to inclination.

Collectively, the optimal geometry 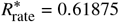 represents a more suitable alternative for practical use, introducing a new design criterion—reducing mechanical load—into the conventional design philosophy that has focused primarily on impedance stability. It provides a concrete design guideline for enhancing comfort through the electrode tip itself, complementing other strategies such as improvements to the electrode holder mechanism.

### 6.2 Limitations and Future Research

This study examined the relationship between electrode tip geometry and SED under specific conditions using static FEM analysis. While the obtained results are valuable, interpreting and applying them requires consideration of the limitations of the modeling approach and evaluation metrics, suggesting directions for future research.

#### 6.2.1 Limitations in Model Simplification

To simplify the analysis, several idealizations were introduced in the model considered in this study.

First, this study primarily focused on a rotation-induced contact scenario, wherein inclination develops during pressing. While this condition is realistic, particularly in applications involving flexible caps, other scenarios—such as initial electrode misalignment or posture changes during use—are also relevant in practice[18]. Considering this, although rotation-induced contact was treated as the representative mechanism, an additional analysis of initially inclined pressing was conducted, with results provided in the Supplementary Material. To establish more comprehensive and versatile design guidelines, future research should incorporate integrated evaluations across multiple contact scenarios.

Second, a single-prong model was adopted to represent the prong geometry. Although the intent was to isolate the influence of tip geometry, this simplification does not capture effects specific to multi-prong configurations, such as mechanical interference between prongs and conformance to scalp curvature—factors that are relevant in practical electrode designs [16,23,27,66].

Third, the scalp was modeled as a single linear elastic body. While this provides a valid first approximation for assessing stress concentration trends associated with tip geometry, greater quantitative accuracy requires extending the model to incorporate the skin’s multilayered structure and nonlinear hyperelastic behavior.

#### 6.2.2 Limitations in the Relationship Between Evaluation Metrics and Practicality

Despite advances in simulation-based design, challenges remain in developing evaluation metrics that effectively bridge simulation results and practical performance.

In this study, SED was used as a mechanical proxy for comfort, based on prior research indicating a correlation between SED and neural responses. However, most of this research has focused on glabrous (non–hairy) skin, such as the fingertips, and the perceptual characteristics of the scalp—where hair follicles are present—may not correspond directly [33,35,39]. To validate the correlation between SED and scalp comfort, future studies should compare subjective comfort ratings from human participants with the results of mechanical analysis.

Additionally, this study did not evaluate electrical contact quality. A geometry optimized for comfort does not necessarily ensure stable contact impedance or a high SNR [10,11,14]. Achieving both comfort and electrical reliability in electrode design will require a combined optimization strategy that integrates mechanical simulation with evaluation of electrical properties.

## 7 Conclusion

This study employed FE analysis to investigate how the tip geometry of dry EEG electrodes influences mechanical loading on the scalp, ultimately aiming to identify a geometry that minimizes this load.

Our findings revealed that while the mainstream hemispherical geometry offers stable performance under tilt, it consistently imposes a high mechanical load on the scalp. In contrast, an intermediate fillet geometry (*R*_rate_ ≈ 0.62) was found to maintain low SED in both normal and tilted conditions. This suggests the potential to design electrodes that balance the tilt stability of hemispherical geometries with the low mechanical load of flat designs, thereby reducing stress on the scalp even under tilted contact.

The geometry ratio identified here represents an optimal solution derived under specific analytical conditions. It provides a mechanical resolution to the trade-off between the “response stability to tilt” offered by hemispherical geometries and the “low initial load” characteristic of flat designs. In other words, it enables the design of electrodes that suppress local scalp loading, even under the practically unavoidable condition of tilted contact, and may form the foundation for a new direction in dry electrode design.

## Acknowledgment

This work was supported by JSPS KAKENHI Grant Number JP22K04014.

