## Supplementary materials for "Design Guidelines for Dry Electrode Tip Geometry in Electroencephalography Measurements: A Proposal Based on the Scalp’s Mechanical Response"

### Supplementary Material for "Design Guidelines for the Tip Geometry of Dry Electrodes for EEG Measurement: A Proposal Based on the Mechanical Response of the Scalp"

*Shunya Araki[1], Shintaro Nakatani[1], and Nozomu Araki[2]*

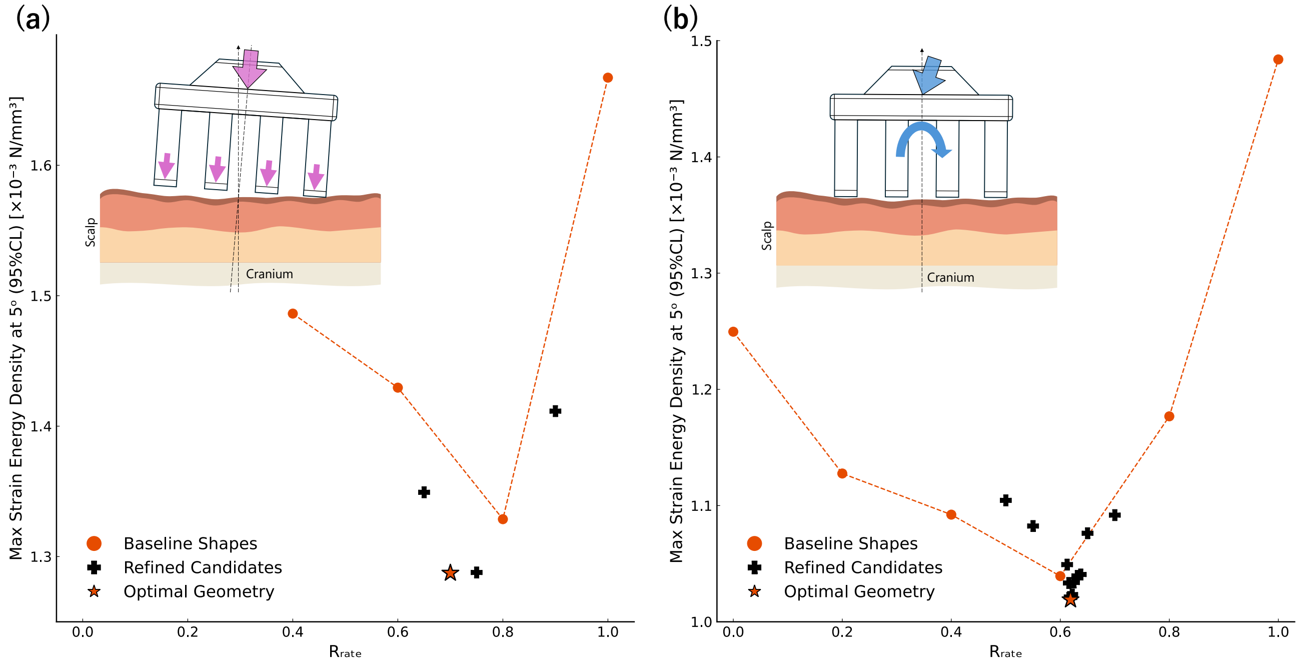


Fig. S1: Comparison of optimal shapes under two different tilted contact scenarios. Both panels show the relationship between $\mathrm{SED}_{\max}$ and the geometry ratio $R_{\mathrm{rate}}$ under a 5° inclination, along with the optimization results from an iterative search. (a) Results for the "initially-inclined pressing" scenario, where the electrode is pressed while already tilted from its initial placement. The optimal geometry was found at $R^*_{\mathrm{rate}} = 0.7$. (b) Results for the "rotation-induced contact" scenario (the main analysis in this paper), where tilt is induced by a non-normal force on the entire electrode. The optimal geometry was found at $R^*_{\mathrm{rate}} = 0.61875$.

As supplementary information, Fig. S1 presents the results of an additional analysis conducted on "initially-inclined pressing," a scenario different from the "rotation-induced contact" that was the primary focus of the main paper. Using the same finite element model as in the main study, a nonlinear static analysis was performed by pressing the prong from an inclined state to achieve a displacement of 0.1 mm at the tip's center. Six types of prong tip geometries were defined using the shape ratio $R_{\mathrm{rate}}$, consistent with the main paper. The analysis revealed that for geometries with $R_{\mathrm{rate}}$= 0.0 and 0.2, the simulation failed to converge due to excessive element deformation at the edges. This result implies the occurrence of extremely high stress concentrations, strongly suggesting that these geometries would involve excessive mechanical loading and a risk of skin damage in practical use. For the other geometries where the simulation converged, the relationship between $\mathrm{SED}_{\max}$ and $R_{\mathrm{rate}}$ exhibited a downwardly convex trend, similar to the findings in the main paper. Therefore, an iterative search was conducted to identify the optimal geometry in this contact scenario, using the same algorithm (i.e., identical learning rate, update rule, and convergence criterion) as in the main study. After four additional analyses, the geometry that minimized the maximum strain energy density was found to be $R^*_{\mathrm{rate}} = 0.7$. This finding indicates that even under a different mechanical mechanism from the rotation-induced contact (optimal solution: $R^*_{\mathrm{rate}} \approx 0.62$), an intermediate fillet geometry (optimal solution: $R^*_{\mathrm{rate}} = 0.7$) similarly minimizes the mechanical load. In this scenario, where the edge penetrates the skin from the initial moment of contact, the more rounded geometry ($R^*_{\mathrm{rate}} = 0.7$) was likely optimal because it was more effective at mitigating the initial stress concentration. This fact strongly supports the robustness of the main conclusion of this paper: that an intermediate fillet geometry with an $R^*_{\mathrm{rate}}$ in the range of 0.6 to 0.7 can consistently reduce mechanical load under various tilted contact conditions, not limited to a specific scenario.



1. Graduate School of Sustainability Science, Tottori University

2. Department of Mechanical Engineering, Graduate School of Engineering, University of Hyogo
